# Machine learning detection of SARS-CoV-2 high-risk variants

**DOI:** 10.1101/2023.04.19.537460

**Authors:** Lun Li, Cuiping Li, Na Li, Dong Zou, Wenming Zhao, Yongbiao Xue, Zhang Zhang, Yiming Bao, Shuhui Song

## Abstract

The severe acute respiratory syndrome coronavirus 2 (SARS-CoV-2) has evolved many high-risk variants, resulting in repeated COVID-19 waves of pandemic during the past years. Therefore, accurate early-warning of high-risk variants is vital for epidemic prevention and control. Here we construct a machine learning model to predict high-risk variants of SARS-CoV-2 by LightGBM algorithm based on several important haplotype network features. As demonstrated on a series of different retrospective testing datasets, our model achieves accurate prediction of all variants of concern (VOC) and most variants of interest (AUC=0.96). Prediction based on the latest sequences shows that the newly emerging lineage BA.5 has the highest risk score and spreads rapidly to become a major epidemic lineage in multiple countries, suggesting that BA.5 bears great potential to be a VOC. In sum, our machine learning model is capable to early predict high-risk variants soon after their emergence, thus greatly improving public health preparedness against the evolving virus.

## Introduction

The COVID-19 pandemic has been characterized by repeated waves of cases driven by the emergence of multiple high-risk SARS-CoV-2 variants, which show increased ability of transmission and evasion of existing immunity by infection or vaccine[1]. Thus, rapidly and early identifying such variants and accurately forecasting their dynamics as they emerge are critical for guiding outbreak response. To effectively prevent and control the COVID-19 epidemic, several high-risk SARS-CoV-2 variants, such as variants of concern (VOC) and variants of interest (VOI), have been designated by the World Health Organization (WHO) based on shared attributes and characteristics that may require public health action (https://www.who.int/en/activities/tracking-SARS-CoV-2-variants). Besides, multiple variants under monitoring (VUM) and VOC lineages under monitoring (VOC-LUM) are designated to investigate if they need prioritized attention and monitoring. Although these variants are monitored by combining epidemiological investigations, genetic sequence-based surveillance and laboratory studies, early prediction is severely lagging behind the emergence of virus strains to warning, which can take as long as 1∼7 months. Computationally, sensitive and accurate early warning methods are urgently needed.

Recently, several studies on evaluation and prediction of SARS-CoV-2 high-risk mutations or variants have been performed using different machine learning models[2-5]. These studies differ in mutations used that are derived from whole genome or specific gene. Based on whole genome mutations, methods for predicting mutation spread[2], PyR_0_[3], and VarEPS[4] were proposed via logistic regression model, Pyro, and random forest, respectively. In detail, methods for predicting mutation spread characterized the potential risks of the single-position substitutions using a logistic regression model, and forecasted the driver mutations for future SARS-CoV-2 VOC according to epidemiology, evolution, and immunology features[2]. PyR_0_ identified numerous mutations and gene regions that will likely increase SARS-CoV-2 fitness using Pyro, and also inferred and ranked lineage fitness by statistical significance[6]. VarEPS showed an evaluation and pre-warning for all known mutations and virtual mutations based on genomics and structural biology characteristics[4], evaluated the risk level of variants and grouped strains by their transmissibility and affinity to neutralizing antibodies using random foreast[4]. Additionally, several studies proposed methods based on mutations specifically in the spike gene[5, 7], e.g. Early Warning System (EWS)[5] could accurately rank SARS-CoV-2 variants for immune escape and fitness potential by combining spike protein structure modelling and large protein transformer language model on spike protein sequences. However, most of these studies considered only those traditional sequence features in a static state, but ignored the evolutionary relationship between variants.

Haplotype network is a common way for illustrating phylogenic results, which not only provides the evolutionary relationship of sequences[8], but also reflects the transmission characteristics of different sequences[9, 10]. A variety of network features in the field of graph theory, such as node degree, betweenness, and network density, can be exploited to study the transmission dynamics[11]. Understanding the role of haplotype network characteristics, and how they are related to virus transmissibility are thus of great importance for performing risk evaluation and prediction of SARS-CoV-2 at variant level. In this work, we comprehensively analyzed the characteristics of SARS-CoV-2 haplotype networks and revealed several important features which are informative for predicting high-risk variants at haplotype level. We further trained a supervised machine learning model (HiRisk-Predictor) by combining all these network features using the LightGBM algorithm[12]. By HiRisk-Predictor, we obtained early prediction with high accuracy on retrospective studies. Prediction based on sequences as of June 15, 2022 showed several emerging high-risk variants, including BA.5 and BA.4 etc. Taken together, our model is able to predict high-risk variants soon after their emergence and thus can improve public health preparedness against ongoing the evolving virus, which could also be applied to the monitoring of any rapidly evolving pathogens with sufficient genomic data.

## Results

### Analysis of haplotype network features for SARS-CoV-2 variants

Haplotype network is fundamental and useful for tracing and visualizing the genealogies of SARS-CoV-2 genomes[13], but network features reflecting virus characteristics of propagation or transmission have not been fully recognized. Here, we find some network features, e.g., node size and node out-degree, may be associated with virus transmission based on a series of snapshots of early data for the Delta variants (**Fig. 1a**). Then we comprehensively analyze the characteristics of haplotype networks, including node size, network centrality (node out-degree and betweenness), sequence growth ratio, geographic information entropy (GIE), and mutation scores in the spike protein (**Fig. 1b**). The distribution of node size (the number of genome sequences in a node) is strongly skewed and follows a power-law distribution y∼x^-2.1^, suggesting that almost all non-observed haplotypes will be small, nevertheless those haplotypes with large size may be major epidemic viruses globally or regionally. The node out-degree (the total number of edges attached to a node) and betweenness centrality (the number of the shortest paths that pass through a node) are indicators of node influence, which exhibit a power law tail distribution (scale-free and robust network) with exponents about 2.29 and 1.75, respectively. This suggests that nodes with large out-degree or betweenness centrality always have great ability to evolve and transmit, and play a dominant role in the evolutionary process of SARS-CoV-2[14]. In all, these two features could help to identify haplotypes who play a “bridge spanning” role in a network.

**Fig. 1.**
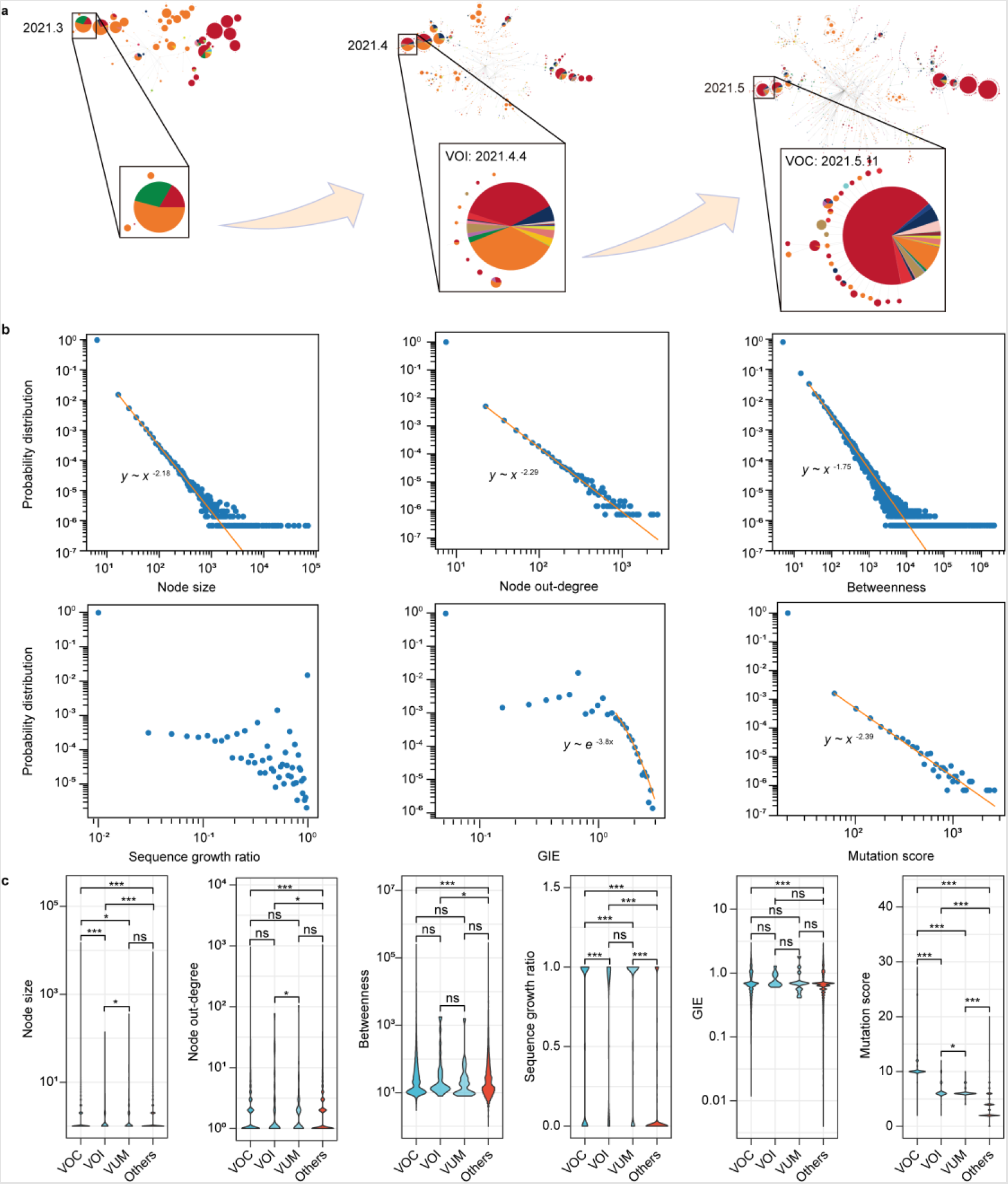
Haplotype network features of SARS-CoV-2 variants. (a) A schematic diagram of haplotype network evolution, with changes for some network features, e.g. node size and geographical transmission range. (b) Double-logarithmic plots of probability distributions of the six haplotype network features, including node size, node out-degree, betweenness centrality, sequence growth ratio, geographic information entropy (GIE), and mutation score in the spike protein. At the tail of probability distributions, orange lines indicate power-law decays and an exponential decay, respectively. (c) Violin plot for the distribution of the features above between WHO current and former VOC, VOI, VUM, and the other variants. The *P* values of Wilcoxon test between any two groups are labeled as * *P* < 0.05, ** *P* < 0.01obvious, *** *P* < 0.001, ns *P* > 0.05.

To quantify the spatiotemporal characteristics of all haplotypes, we calculate the sequence growth rate of a haplotype node and the geographic information entropy (GIE) representing the geographical transmission range. From the distribution of sequence growth rate, we find that 98% of the haplotypes grow slowly (sequence growth rate < 0.01), while a small number of haplotypes increase rapidly (sequence growth rate > 0.9) that need to be concerned. By analyzing GIE, we also observe that 96% of the haplotypes are locally propagated variants (GIE < 0.05). The exponential decay of GIE distribution tail suggests that a few haplotypes (0.5%) with large GIE (> 1.0) should be concerned due to their transmissibility to a wider geographical area. As mutations in the spike protein may affect the transmission of the virus by changing the binding stability with the host cell receptor angiotensin-converting enzyme 2 (ACE2) and the antibody affinity of SARS-CoV-2[15], a mutation score for each haplotype is introduced to quantitatively evaluate the strength of the binding stability and antibody affinity. The probability of haplotypes with high mutation scores is small, yet these haplotypes may be high-risk variants. Collectively, it is necessary to integrate all these features to predict the risk of SARS-CoV-2 variants.

By comparing the distributions of these network features in VOC, VOI, VUM, and the other haplotypes, significant differences (Wilcoxon test, *P* < 0.05) are observed between high-risk variants (VOC and VOI) and the others except for GIE (**Fig. 1c**). While between VUM and the others, only two features, the sequence growth ratio and mutation scores in the spike protein, are significantly different and informative. Together, our results reveal that those nodes with large node size or out-degree, and nodes with high growth rate, GIE or mutation score are more likely to be high-risk variants and these haplotype network features have potential to predict high-risk variants.

### An accurate machine learning model for high-risk variants prediction

To predict high-risk variants, we construct a machine learning model using the features mentioned above (**Supplementary Fig. 1a**). Focusing on the growing or emerging variants in a time period, potential high-risk haplotypes are defined as those which satisfy the following empirical criterions: node size (number of sequences >1), node out-degree > 0, GIE > 0, and sequence growth ratio > 0.75. To further characterize the connectedness of potential high-risk haplotypes (the number of potential high-risk haplotypes in a connected subnetwork) in a haplotype network, we introduce a new feature called connectivity of nodes. Then, we construct a training dataset consisting of 574 potential high-risk haplotypes calculated from SARS-CoV-2 sequences with collection date up to October 31, 2020, November 30, 2020, and December 31, 2020, where all haplotypes belong to Alpha and Beta variants are labeled as positive samples (90, 15.7%), while the others are labeled as negative samples (484, 84.3%). We also construct a series of testing datasets including potential high-risk haplotypes. In these testing datasets, those potential high-risk haplotypes are labeled as high-risk haplotypes when they are known VOC or VOI defined by WHO.

Next, we determine which features are significant for predicting high-risk haplotypes. When we compare the distribution of those haplotype network features between positive and negative samples, significant differences (Wilcoxon test, *P*<0.05) are observed for all features (**Fig. 2a**). Among them, the two most significant (*P*<0.001) features are mutation scores in the spike protein and connectivity of nodes. However, none of these features alone could classify the high-risk variants accurately. When we reduce the dimension of features into two-dimensional space by t-Distributed Stochastic Neighbor Embedding (t-SNE, setting perplexity as 50 and iteration as 1000)[16], clear boundaries between positive and negative samples are observed (**Fig. 2b**), suggesting that high-risk variants can be predicted by integrating all these network features.

**Fig. 2.**
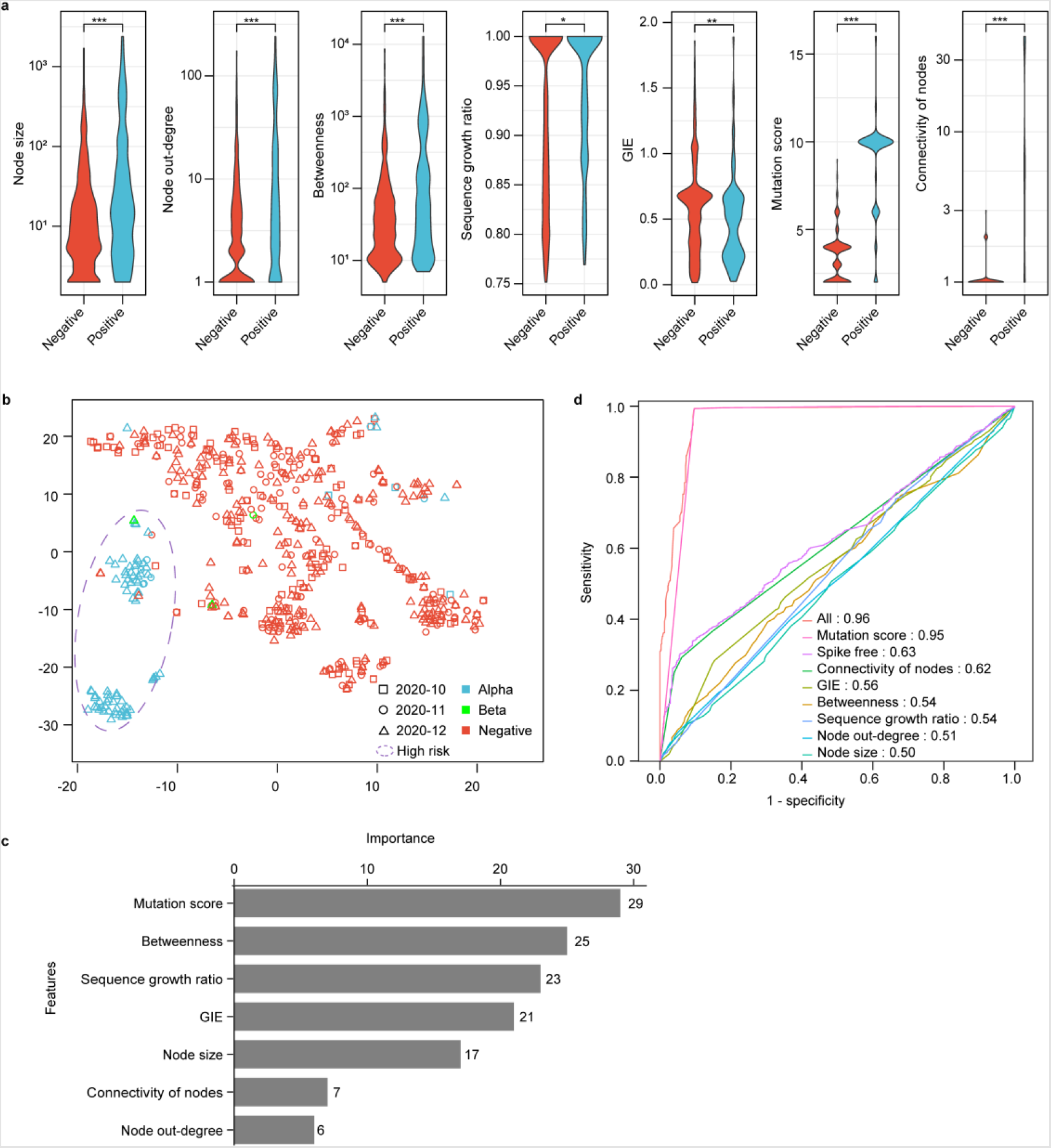
Performance evaluation of haplotype features for high-risk variants prediction. (a) Violin plot for haplotype network features in the training dataset, including node size, node out-degree, betweenness centrality, sequence growth ratio, geographic information entropy (GIE), connectivity of nodes, and mutation scores in the spike protein. “Positive” represents positive samples which are Alpha and Beta haplotypes and others are labelled as “Negative”. Significance test between positive and negative samples were estimated by Wilcoxon test (* *P* < 0.05, ** *P* < 0.01, *** *P* < 0.001). (b) Haplotypes illustration in two-dimensional space by t-SNE (setting perplexity as 50, iteration as 1000). (c) The importance of network features in HiRisk-Predictor model. (d) Performance of prediction models using different features.

We then train a supervised machine learning model (HiRisk-Predictor) by combining all these network features using the LightGBM algorithm. The importance of these seven features for the training model is shown in **Fig. 2c**, and the mutation score in the spike protein has the greatest impact on the prediction of risk of SARS-CoV-2 variants. Moreover, HiRisk-Predictor shows remarkable discrimination between high-risk and low-risk haplotypes on the testing datasets, reflected by a high AUC value of 0.96 (**Fig. 2d**). To determine the diagnostic ability of each individual feature, we train the model using individual features. AUC of the seven models shows that the mutation score model has the best diagnostic ability, and HiRisk-Predictor combining all seven features performs better than any models with individual features (**Fig. 2d**).

### Early and accurate detection of high-risk variants on retrospective datasets

We perform retrospective prediction on a series of different testing datasets generated from SARS-CoV-2 sequences with collection date up to different dates ranging from January 2021 to June 2022. To identify high-risk haplotypes, we define a risk score which is the probability of a haplotype to be positive in HiRisk-Predictor. The predicted risk score for all potential haplotypes based on collection date of sequences (all sequence data available at the time of analysis) and submission date (sequence data available until that time) are available on RCoV19 (https://ngdc.cncb.ac.cn/ncov/monitoring/risk). If the predicted risk score of a haplotype is greater than 0.5, it will be a high-risk haplotype. In all predicted high-risk haplotypes, there are not only haplotypes of Alpha and Beta, but also haplotypes of other current and former VOC or VOI variants (e.g., Delta and Omicron). This indicates that HiRisk-Predictor learns well not only the patterns of Alpha and Beta from training datasets, but also the patterns of other high-risk variants. When based on collection date of sequence, the detection date for eight VOC and VOI variants (e.g., Delta, Iota, Kappa, Lambda, and Mu) is several months (one to four months) earlier than WHO announced (**Fig. 3a**). Taking Delta as an example, seven haplotypes have been detected to be high-risk variants on January 10, 2021, while Delta was officially designated as VOI on April 4, 2021 and VOC on May 11, 2021 by WHO. While for Omicron, our detection date by HiRisk-Predictor is equivalent to WHO-announced date. This is because Omicron raised global concerns immediately as it harbors a large number of mutations including more than 30 changes in the spike protein and 15 in the receptor-binding domain (RBD) of spike[17]. However, our model does not predict Theta and Zeta as high-risk variants, suggesting that HiRisk-Predictor is not sensitive to some VOI variants. As known from these analyses, HiRisk-Predictor is able to predict all VOC and most VOI variants before they become a worldwide epidemic. However, when we predict based on submission date of sequence, the detection date of most VOC and VOI variants lag behind WHO-announced date by one to three weeks except for Mu. This indicates that sequence data sharing timely is important for early warning and pandemic prevention. As reported, the median collection to submission time (CST) lag is from one day to over one year for SARS-CoV-2 sequences to GISAID[18], so that we suggest performing efficient surveillance and pre-warning based on both collection and submission dates of sequences for the needs of practical application.

**Fig. 3.**
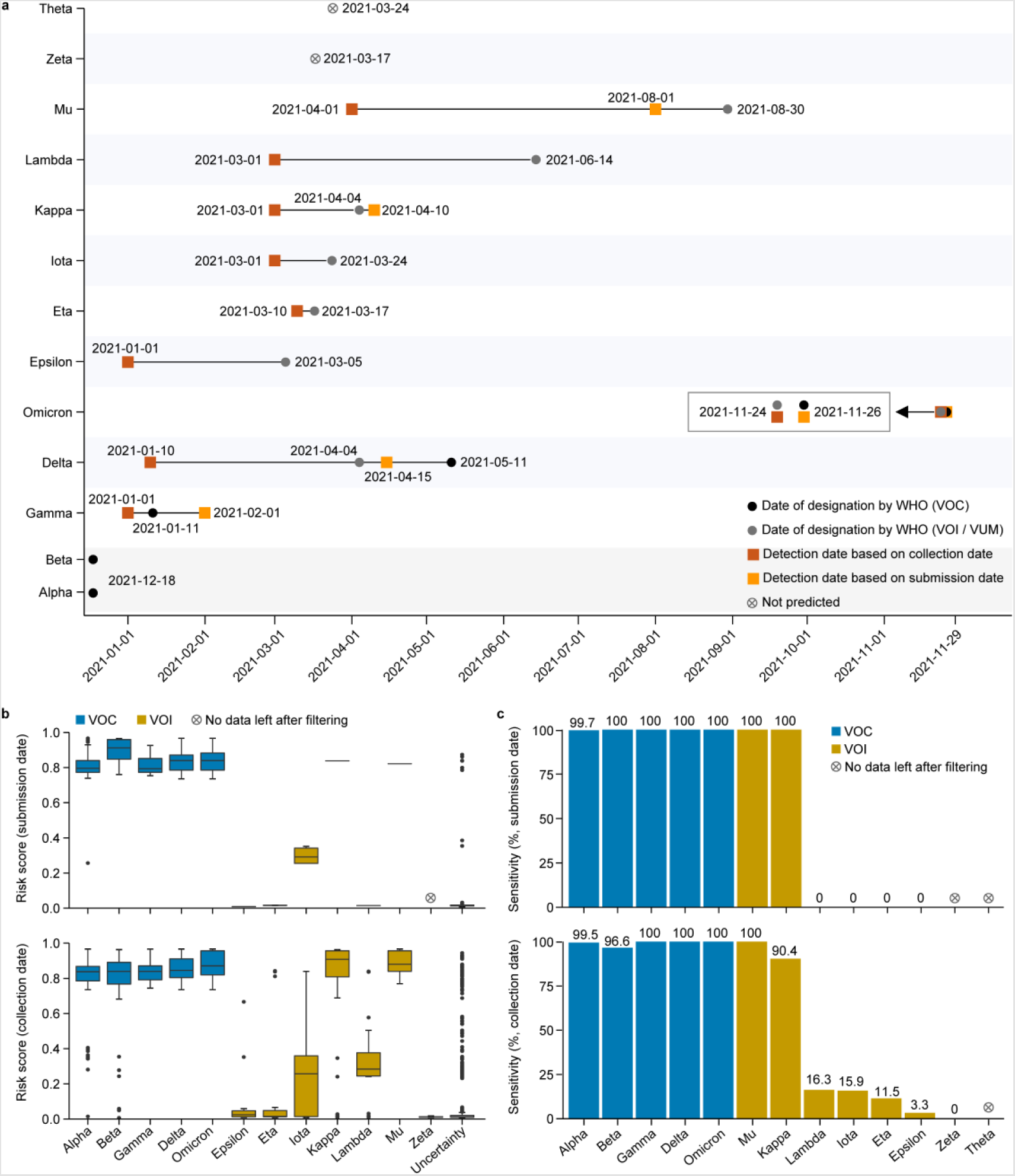
Retrospective prediction results on a series of different testing datasets. (a) Detection dates for VOC and VOI variants based on sample collection and sequence submission date, which indicated as red and orange rectangle, respectively. Dates of designation by WHO are indicated in black and gray circle, respectively. (b) Distribution of predicted risk score for high-risk haplotypes labeled to VOC and VOI variants. (c) The sensitivity (proportion of predicted high-risk haplotypes to potential high-risk haplotypes) for each VOC and VOI variant. The top and bottom panels were predicted by submission and collection date, respectively, in (b) and (c).

Can our predicting model distinguish between VOC and VOI variants? As each haplotype is given a predicted quantitative risk score, we compare the predicted median risk score of VOC and VOI variants, and find that most VOC variants (e.g., Beta, Delta, and Omicron) have risk scores greater than 0.8, which are higher than those for VOI (e.g., Epsilon and Iota) and other low risk variants (**Fig. 3b**). Additionally, based on the number distribution of predicted high-risk haplotypes on a series of different testing datasets, we notice that the number of high-impact variants (e.g., Alpha, Delta, and Omicron) is higher than those of low-impact ones such as Gamma, Kappa, and Mu (**Supplementary Fig. 2**). By calculating the sensitivity for each WHO label, the VOCs’ (> 0.9) are much greater than VOIs’ (**Fig. 3c**). According to these observations, HiRisk-Predictor could detect VOC variants accurately based on publicly available sequence data and thus is suitable for routinely surveillance.

### Another potential VOC variant: BA.5 lineage

With publicly available sequences as of January 15, 2022, HiRisk-Predictor detects 224 high-risk haplotypes which belong to 15 Pangolin lineages[19] (**Fig. 4a**). The median risk scores for these lineages are greater than 0.7. Among these lineages, BA.5, BA.4, BA.2.13, and BA.2.12.1, designated recently as VOC-LUM variants by WHO (https://www.who.int/en/activities/tracking-SARS-CoV-2-variants), have higher risk scores. A new study reported that the effective reproduction numbers of these lineages are greater than that of the original BA.2, due to the acquisition of a mutation at the L452 residue of the spike protein[20]. Especially, BA.5, possessing the highest risk score among these lineages, has started to grow in frequency during the past weeks as known from the prevalence of lineages (**Fig. 4b**). When we compare the percentage of the total daily Omicron infections since its emergence, we also observe that the infections of BA.5 raise to ∼40% within approximate 150 days, which is lower than BA.2 but higher than other high-risk lineages (**Fig. 4c**). Comparing with these lineages, the high transmissibility and growth advantage of BA.5 could be enhanced by stronger neutralization evasion due to the D405N and F486V mutations[21].

**Fig. 4.**
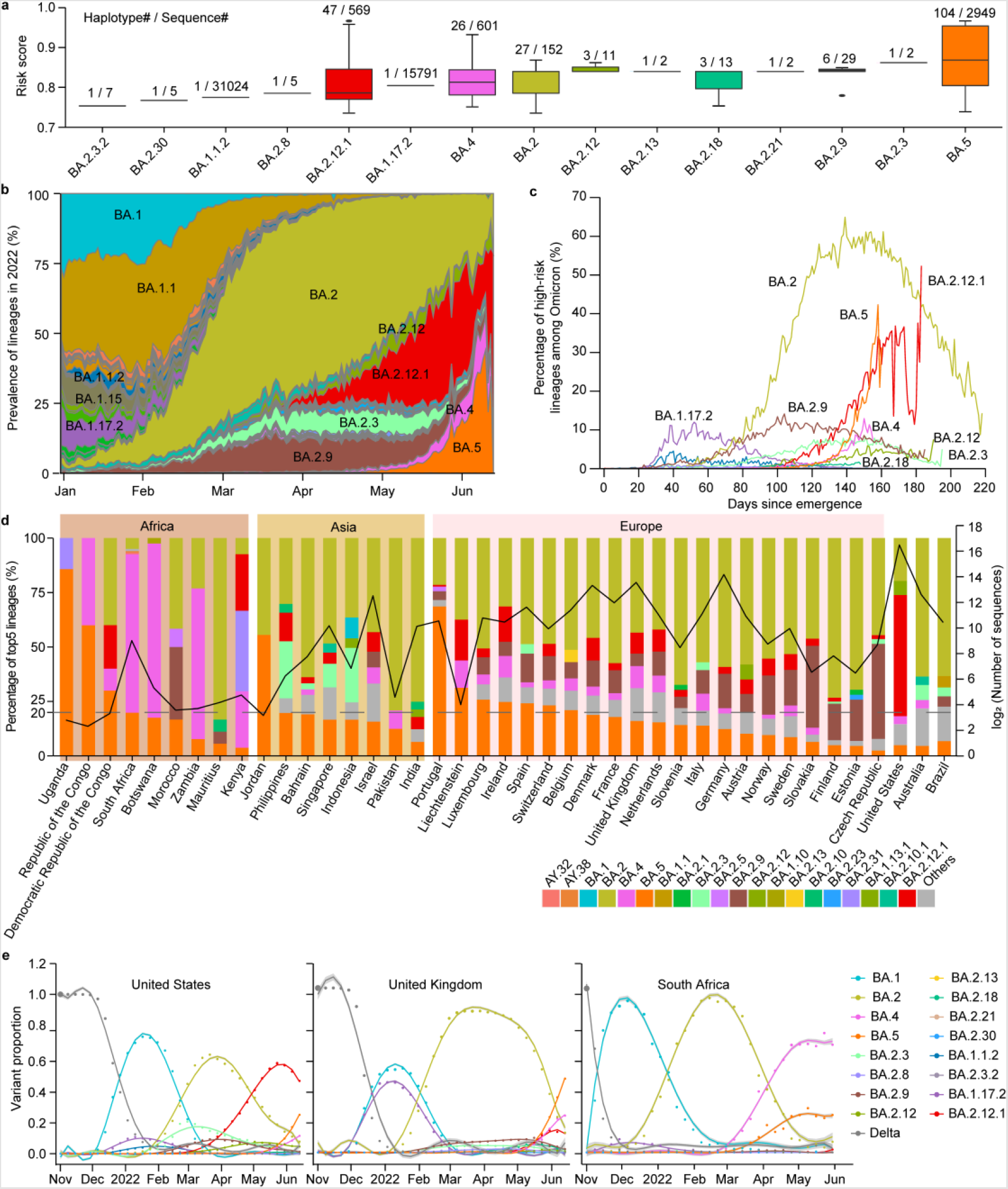
High-risk variants prediction based on released sequences as of June 22, 2022. (a) Risk score distribution of predicted high-risk haplotypes organized according to Pangolin lineages. (b) Genomic prevalence of lineages from January 2022 to June 2022. (c) Percentage of high-risk lineages among all Omicron sequences since its emergence. (d) Percentage of five predominant lineages (based on released sequences collected from May 15, 2022 to June 15, 2022) in countries with prevalent of BA.5. (e) Epidemic dynamics of SARS-CoV-2 lineages in USA, UK, and South Africa. The results for weekly sequence frequency and modelled generalized linear proportions (based on a multinomial logistic regression) of high-risk lineages are shown. BA.5 is growing and competing with BA.4, BA.2.12.1, and BA.2.13.

At the time of this writing, BA.5 has been reported in 69 countries and regions (**Fig. 4d, Supplementary Fig. 3a**). It becomes dominant (portion of sampling sequences > 25%) in Uganda, Republic of the Congo, Democratic Republic of the Congo, Jordan, Portugal, etc., according to sampling sequences in the recent month (**Fig. 4d**), and is spreading rapidly in Luxembourg, Portugal, Ireland, etc., since its emergence (**Supplementary Fig. 3b**). As SARS-CoV-2 genomic surveillance and sequence availability varies across countries[22], we compare the proportions of lineages since November 2021 in Portugal; one of the five countries with BA.5 dominancy, BA.5 is swiftly replacing the fifth wave of BA.2 (**Supplementary Fig. 3c**). While in South Africa, USA, and UK with high level of routine genomic surveillance and high availability of SARS-CoV-2 sequences, BA.5 is growing and competing with BA.4, BA.2.12.1, and BA.2.13 (**Fig. 4e**). It is inferred that BA.5 may go on spreading to more countries and regions. While the size of the BA.5 wave may vary from place to place which depends on their immunity profile[23]. Altogether, the possibility of a new wave of COVID-19 epidemic around the world should not be ignored.

## Discussions

Here we constructed a machine learning model (HiRisk-Predictor) that is capable of accurately identifying SARS-CoV-2 high-risk variants via LightGBM. HiRisk-Predictor detected Omicron variants with high risk score and a large number of high-risk haplotypes at the beginning of their emergence. Sensitivity to Omicron may be due to its larger number of mutations accumulated in potential mouse origins[24] or its increased transmission efficiency[25]. For other high-risk variants, the detection date of HiRisk-Predictor for VOC and most VOI variants is earlier than that of WHO, but a delay from the date of emergence to the detection date still exists. The delay may be caused by the following three factors: first, more features reflecting immunity and epidemiology, such as Epi Score[2], CD4 response, CD8 response[26], should be introduced in HiRisk-Predictor for improving the sensitivity in the future. Second, the large gap of sequencing rate between low- and middle-income countries and developed countries may affect the accuracy of some features in HiRisk-Predictor, such as node size, GIE, etc.[27] The delay time may come from the recessive transmission of the variant in under-developed countries with limited sequencing and genomic surveillance capabilities, which is critical in monitoring SARS-CoV-2. Third, the collection to submission time (CST) lag[18] may also increase the delay time due to the inability to obtain real-time data. Therefore, improving sequencing rate and submitting genomes to open access platforms are both crucial to develop pre-warning system for SARS-CoV-2 high-risk variants.

As the epidemic evolution, the data of SARS-CoV-2 genomes is growing by tens of thousands every day (https://ngdc.cncb.ac.cn/ncov/). Routinely monitoring and warning are important and helpful in dynamically controlling the epidemic. With the newest sequences (up to July 25, 2022), HiRisk-Predictor detected several emerging variants (**Supplementary Fig. 4**), including BA.5 and its decedent sub-lineages. According to PANGO COMMITTEES, dozens of sub-lineages of BA.5 have been discovered, e.g., BA.5.2, BA.5.2.1 in the past few months (https://cov-lineages.org/lineage_list.html). Moreover, the prevalence of these lineages has predominated among all publicly available sequences at rapidly increasing rates (from about 30% to 60%) (**Fig. 4b** and **Supplementary Fig. 4b**) in just one month. This is in accordance with the prediction by HiRisk-Predictor that BA.5 is a high-risk variant (**Fig. 4a**), convincingly confirming that HiRisk-Predictor is able to accurately monitor and detect high-risk variants of SARS-CoV-2. More importantly, another two divergent BA.5 subvariants, BF.5 and BF.1 (https://cov-lineages.org/lineage_list.html), were detected to have the highest risk scores (**Supplementary Fig. 4a**). According to recent reports from PANGO COMMITTEES and Outbreak.info, BF.5 is locally prevalent, mainly in Israel (65.0%) and USA (9.0%), while BF.1 is widespread, mainly in multiple countries in Europe and USA (https://outbreak.info/situation-reports?pango=BF.1, https://outbreak.info/situation-reports?pango=BF.5, https://cov-lineages.org/lineage_list.html). Hence, these two lineages may be newly-developing VOC variants. To prevent another potential pandemic of SARS-CoV-2, further epidemic statuses of these lineages should be monitored continually.

In summary, it is a new perspective to identify high-risk variants of SARS-CoV-2 by the machine learning model HiRisk-Predictor, trained with both haplotype network features and mutations. High accuracy and sensitivity of HiRisk-Predictor bears promise to improve public health preparedness against ongoing evolving virus, which could be also applied to the monitoring of any rapidly evolving pathogens with sufficient genomic data.

## Methods

### Sequences data and haplotype network construction

All SARS-CoV-2 sequences, genomic variants, and metadata were downloaded from GISAID[28] and RCoV19[29] (as of July 25, 2022). To characterize the diversity of virus sequences, the population mutation frequency (PMF) for each mutated site in a time window (within every month) was calculated. Those non-UTR mutations with PMF > 0.005 were screened out for haplotype network construction by McAN[30].

### Haplotype features

To predict high-risk variants accurately, seven features were extracted for each haplotype in haplotype network, including node size (number of sequences), node out-degree, betweenness, geographic information entropy (GIE), sequence growth ratio, mutation scores, and connectivity of nodes. Haplotype network features (node out-degree, betweenness centrality, connectivity), were calculated using igraph tool[31]. GIE for each haplotype was calculated according to the following formula,

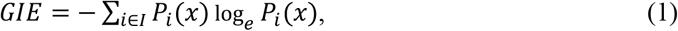

Where *P*_*i*_(*x*) is the probability of a sample *x* in a haplotype at location *i, I* is the set of all locations. Sequence growth ratio of a haplotype was calculated by the following formula:

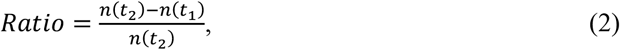

Where *n*(*t*_1_) and *n*(*t*_2_) represent the number of sequences released before time *t*_1_ and *t*_2_ in a haplotype.

Each haplotype is composed of different mutations. Effects of a mutation on binding capacity of ACE2, neutralizing antibodies, and risks of amino acid substitutions were obtained from VarEPS[4]. The mutation score of mutations in spike protein for haplotype *h* is defined to evaluate these effects, as following:

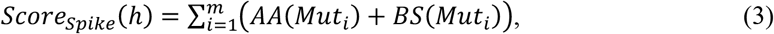

where *Mut*_*i*_, *i*=1, …, *m* are all spike protein SNVs that haplotype *h* has, *AA*(*Mut*) and *BS*(*Mut*) represent the risk level of antibody affinity and binding stability of a single-nucleotide variant (SNV) *Mut*, respectively. All of these parameters are assessed by SAAMBE[32].

Potential high-risk haplotypes were defined as: node size > 1, node out-degree > 0, GIE > 0 and sequence growth ratio > 0.75. For each potential high-risk haplotype, we extracted connectivity feature of nodes, the size of the connected branch of potential high-risk haplotypes. The connectivity was calculated through the connected component of the network only with potential high-risk haplotypes.

Because not all seven features follow normal distributions, we chose Wilcoxon test to calculate *P* values to compare the distribution between any pair of categories by R package and GGsignif[33].

### Benchmark dataset construction and feature extraction for high-risk SARS-CoV-2 variants detection

We constructed a series of training and testing datasets (**Supplementary Fig. 1**). First, individual haplotype network was constructed by McAN[30]. Second, the seven features for each haplotype in haplotype network were extracted. Third, for each potential high-risk haplotype, we assigned positive labels (in the haplotype, more than 80% sequences are VOC or VOI) and negative labels (in the haplotype, more than 80% sequences are neither VOC nor VOI), and unknown for others according to WHO definition. For the training dataset, Alpha and Beta haplotypes were assigned as positive samples, while others as negative. Finally, we trained HiRisk-Predictor using LightGBM. For the testing datasets, all potential high-risk haplotypes with WHO labels are positive samples, while others are negative. To assess the performance of HiRisk-Predictor, the receiver operating characteristic (ROC) curve and the area under ROC curve (AUC) were calculated by R package and pROC[34].

## Acknowledgements

We acknowledge the sample providers and data submitters listed in https://download.cncb.ac.cn/GVM/Coronavirus/prediction/.

## Funding

This work was supported by the National Key Research & Development Program of China (2021YFF0703703, 2021YFC0863300 to S.H.S.), the Strategic Priority Research Program of the Chinese Academy of Sciences (XDB38060100 to Y.M.B.), the National Natural Science Foundation of China (32170678 to W.M.Z., 32270718 to S.H.S.), Youth Innovation Promotion Association of CAS (Y2021038 to S.H.S.), and the Beijing Nova Program (Z211100002121006 to L.L.).

### Conflict of interest statement

None declared.

## Author contributions

Conceptualization, bioinformatics analysis: S.H.S., L.L., C.P.L., and N.L., Writing, original draft: S.H.S., L.L., Web development: D.Z., Review and editing: Y.B.X., Z.Z., Y.M.B., and W.M.Z.

## Notes

### Competing Interest Statement

The authors have declared no competing interest.

