## Supplemental Figures for "Machine learning detection of SARS-CoV-2 high-risk variants"

### Supplementary Figures

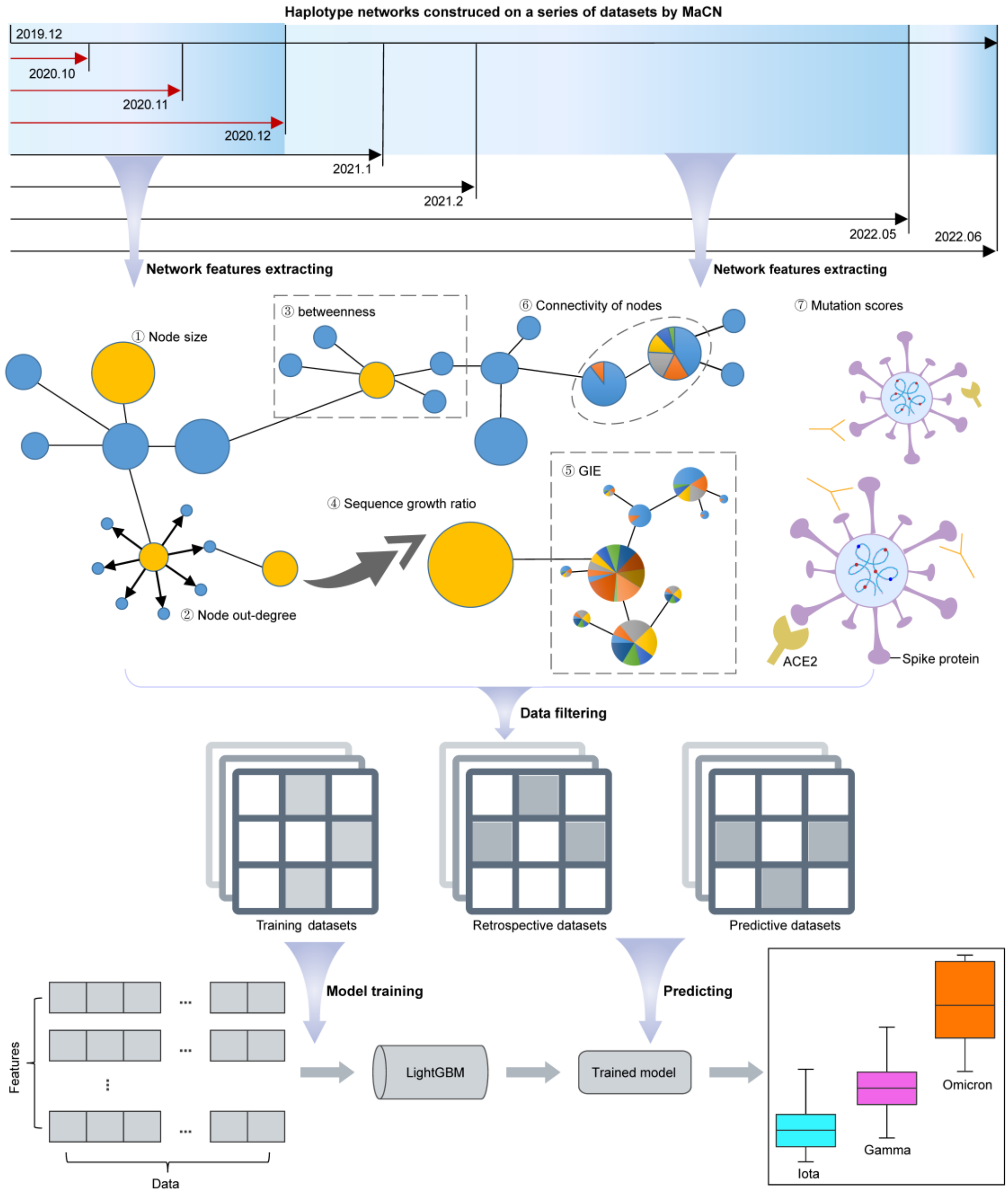


**Supplementary Fig. 1 Schematic illustration of the deep learning-based high-risk variants prediction.**

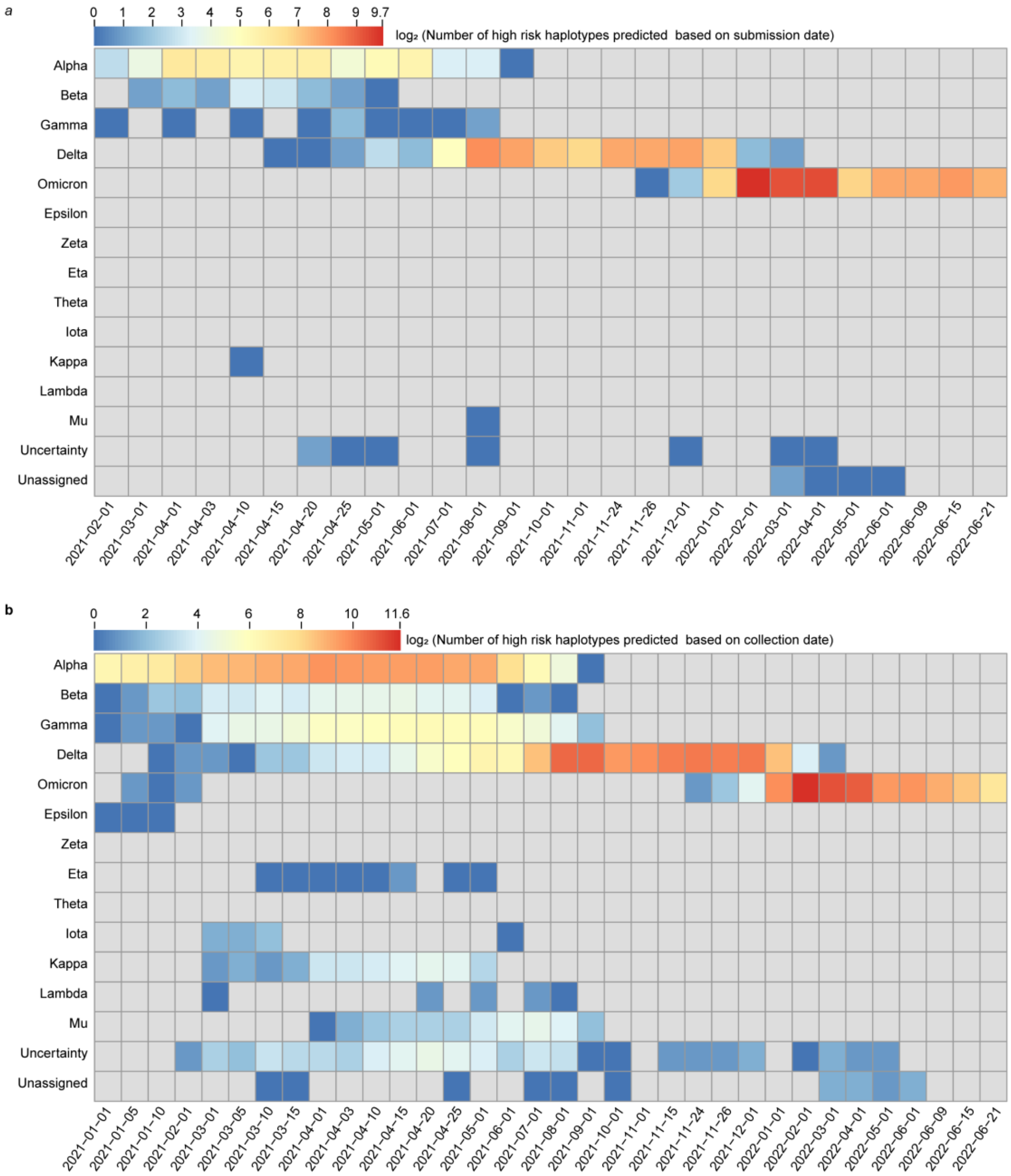


**Supplementary Fig. 2 The number of predicted high-risk variants on testing datasets.**

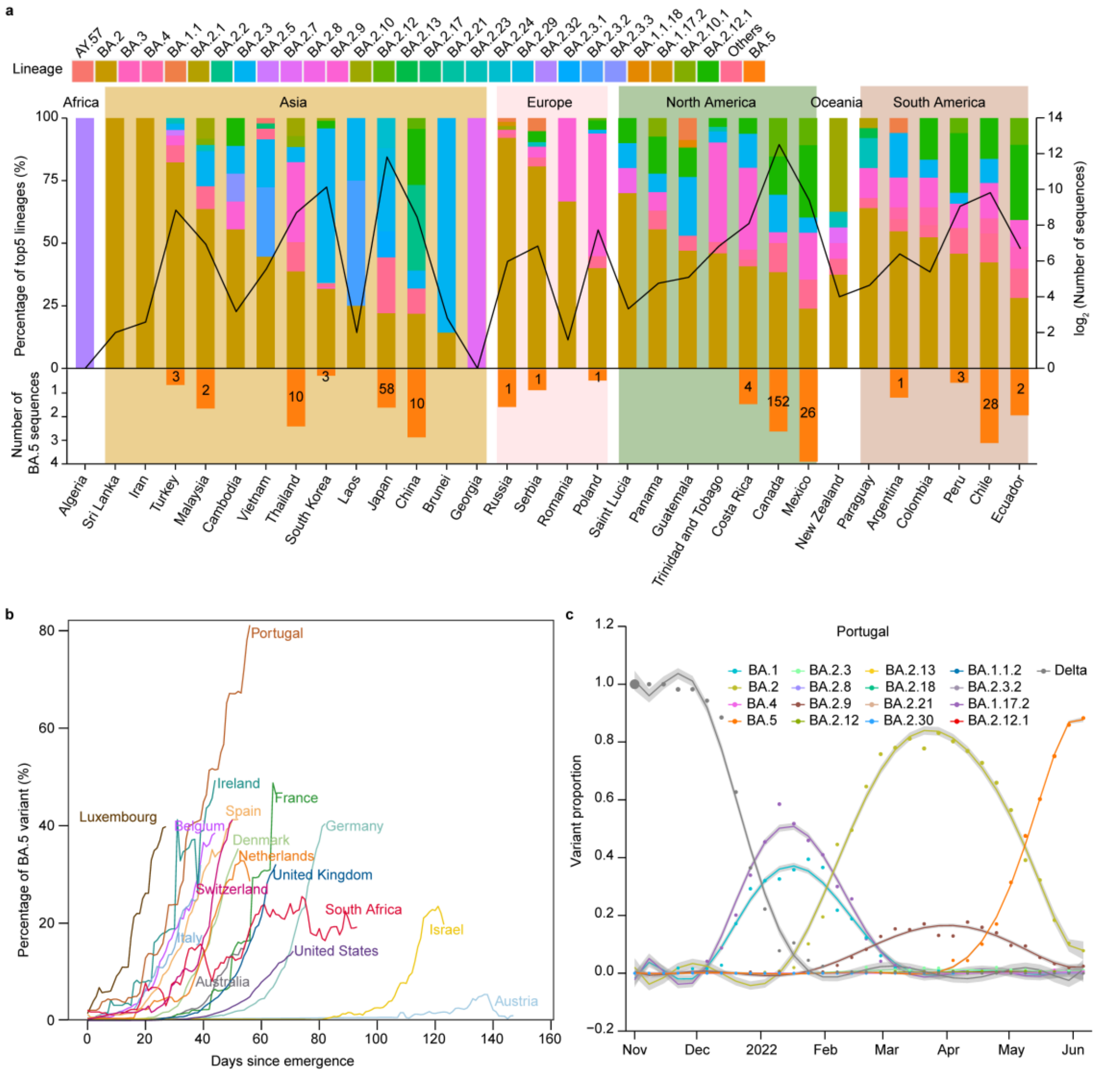


**Supplementary Fig. 3 BA.5 Lineage distribution.**

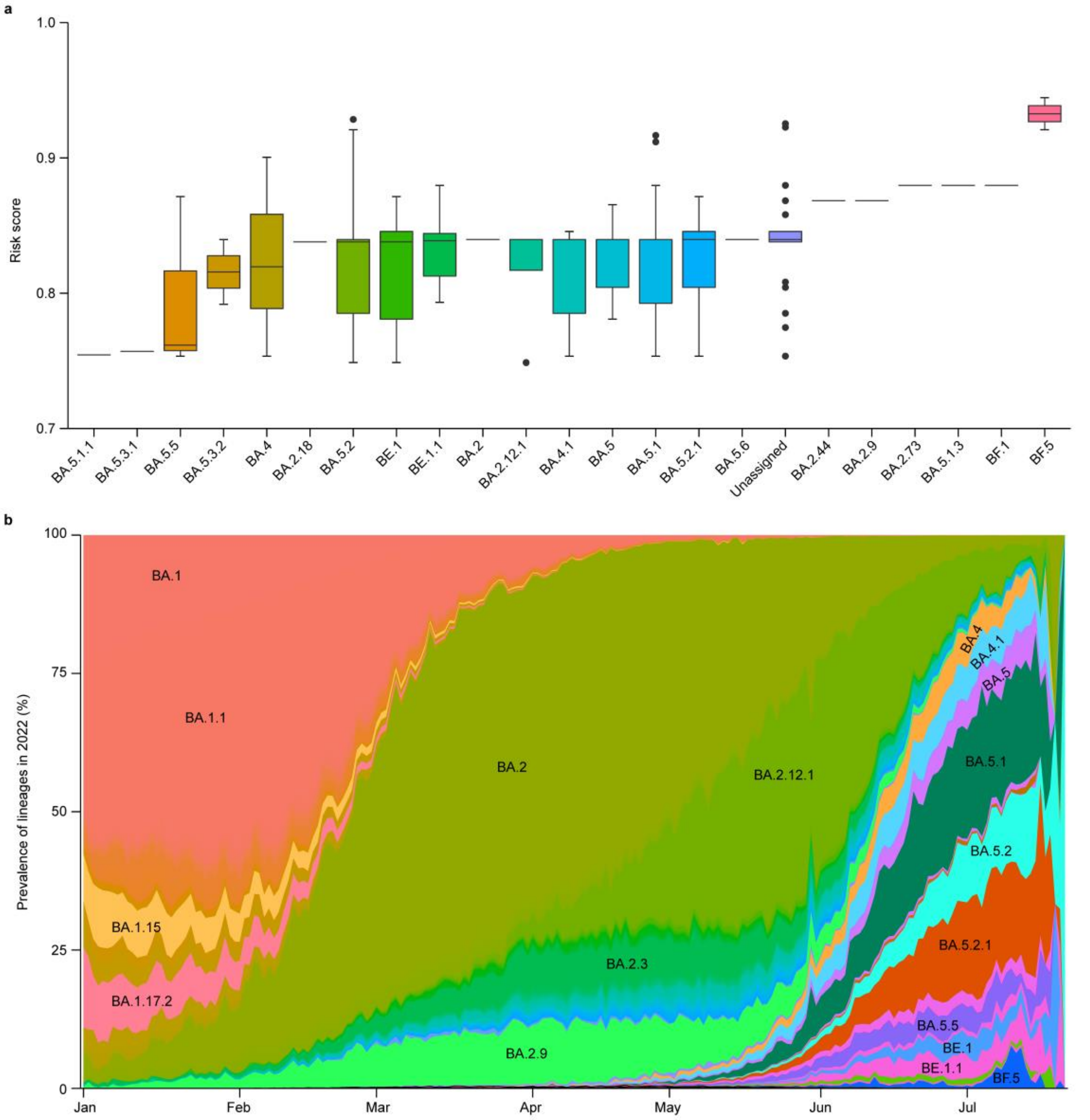


**Supplementary Fig. 4 High-risk variants prediction based on released sequences as of Jul 25, 2022.**
